# Genomic analysis supports Cape Lion population connectivity prior to colonial eradication and extinction

**DOI:** 10.1101/2023.07.31.551389

**Authors:** A. de Flamingh, T.P. Gnoske, A.G. Rivera-Colón, V.A. Simeonovski, J.C. Kerbis Peterhans, N. Yamaguchi, K.E. Witt, J. Catchen, A.L. Roca, R.S. Malhi

## Abstract

Extinct Cape lions (*Panthera leo melanochaitus*) formerly ranged throughout the grassland plains of the “Cape Flats” in what is today known as the Western Cape Province, South Africa. Cape lions were likely eradicated because of overhunting and habitat loss after European colonization. European naturalists originally described Cape lions as “Black-maned lions” and claimed that they were phenotypically distinct. However, other depictions and historical descriptions of lions from the Cape report mixed or light coloration and without black or extensively developed manes. These findings suggest that, rather than forming a distinct population, Cape lions may have had phenotypic and genotypic variation similar to other African lions. Here we investigate extinct Cape lion genome characteristics, population dynamics and genetic distinctiveness prior to their extinction. We generated genomic data from two historic Cape lions to compare to 118 existing high-coverage mitogenomes, and low-coverage nuclear genomes of 53 lions from 13 African countries. We show that, before their eradication, lions from the Cape Flats had diverse mitogenomes and nuclear genomes that clustered with lions from both southern and eastern Africa. Cape lions had high genome-wide heterozygosity and low inbreeding coefficients, indicating that populations in the Cape Flats went extinct so rapidly that genomic effects associated with long-term small population size and isolation were not detectable. Our findings do not support the characterization of Cape lions as phylogeographically distinct, as originally put forth by some European naturalists, but rather highlights how alternative knowledge-systems, e.g., Indigenous perspectives, could potentially further inform interpretations of species’ life histories.

## Introduction

Lions (*Panthera leo* ssp.) formerly ranged across three continents - Africa, Eurasia, and America (Hemmer, 1974; Yamaguchi et al., 2004; de Manuel et al., 2020) - but now only exist as one small population in India and as mostly fragmented and/or isolated populations in Africa (Nowell and Jackson, 1996; Curry et al., 2021; Bertola et al., 2022). The extinct Cape lion formerly ranged throughout the region south of the Orange River (Mazák, 1975), in the various diverse and unique ecological biomes and climactic zones found throughout the ‘Cape’, including the grassland plains of the South African interior. These grasslands are colloquially referred to as the “Cape Flats” and lie west of the Great Escarpment (Mazák, 1975) in what is today known as the Western Cape Province, South Africa. Cape lions were also thought to have occurred in the southern parts of what is known today as the Northern, Eastern and Western Cape Provinces, and the southwest parts of the Free State Province (Burchell, 1824; Smith, 1842; Skead, 1980). The Cape lion population was severely impacted by European settlements and agricultural practices in the mid-1600s (Bryden, 1889; Guggisberg, 1963; Mazák, 1975), as were many African herbivore (Morrison et al., 2007; Ripple et al., 2015) and carnivore populations (Beinart, 1998; Stadler, 2006; Rust and Taylor, 2016). Permanent European colonization, primarily Dutch settlement of the Cape Peninsula beginning in 1652, resulted in Cape lions being hunted as bounty to protect livestock and humans. Documentation of lions being killed are recorded in the journals of the Dutch Governor Jan van Riebeeck (Thom, 1952; Skead, 1980; Stadler, 2006). Cape lions were eventually hunted to extinction in the Western Cape by the 1850s but may have remained until around 1870 in the eastern range of their historical distribution; their complete disappearance was coincidental with the collapse of the ungulate populations in the region (Skead, 1980, 1987).

European naturalists historically described Cape lions as “Black-maned lions” (e.g., Griffith 1821; Smith 1842; Mazák 1964; Mazák 1975). These authors claimed that Cape lions were phenotypically distinct from other lion populations. As one of the earliest Cape lion descriptions, Griffith 1821, from a personal communication from Charles Hamilton Smith, notes that Cape lions were:

*“… a very curious and singular variety… It was nearly the size of the common African lion; but, when compared therewith, was rather thicker altogether, and quite as heavy; the head and muzzle were broader and more pug-shaped; the under jaw was more projecting; the ears larger, slightly acuminated, and black. In character it was more uneasy and restless. The mane was perfectly black, and covered half of the back, and the whole length of the belly*.*”*

In 1842 Smith himself confirms the characteristics noted earlier by Griffith 1821; however, Smith’s publication is considered the ***type*** description since it includes a date, journal and morphological discussion (Smith, 1842):

*“The species is of the largest size, with a bull dog head; the facial line much depressed between the eyes; large pointed ears edged with black; a great mane of the same colour extending beyond the shoulders; a fringe of black hair under the belly; a very stout tail, and the structure in general proportions lower than in other Lions. Habitat the Cape*.*”*

Despite these early descriptions that point towards phenotypic distinctiveness, written accounts also report lions with light and/or manes of mixed dark and light mane coloration (Burchell, 1824; Bennett, 1829). For example, only one of four adult male lion skins that were displayed at the Van Riebeeck Cape Fort Museum Gate, Great Hall and Governor’s Mansion in Cape Town in the 1660s was noted to have a dark mane(van Riebeeck, 1952; Skead, 1980).

Initial support for the Cape lions’ distinctiveness was based on a single, wild caught but captive-raised lion described first in Griffith (1821), then by Smith as a personal communication, and again by Smith (1842) in the formal type description (Supplementary File 1). Rather, Cape lions may have had phenotypic and genotypic variation similar to other African lion populations where manes of mature lions vary, with intermediate types, from black to light yellow (West and Packer, 2002). Such similarity in complete nuclear genomes would align with previous studies that analyzed a short mitochondrial segment and showed that Cape lions do not form a phylogenetically distinct group (Barnett et al., 2006b, 2006a). To understand the long-term evolutionary history of extinct and modern lions, de Manual et al. (2020) investigated the nuclear genomic patterns of 20 lions that occurred across a 30 000-year timespan, two of which were Cape lions. De Manual et al. (2020) found that the genomic diversity of these two Cape lions fall within the diversity of South African lions; however, this analysis considered only 10 historical and 4 extant lions from Africa, and the contextualization of Cape lion nuclear genome characteristics as part of expansive genomics patterns across Africa during historic times has not been investigated.

Here we investigate extinct Cape lion genome characteristics, phylogenomic associations and genetic distinctiveness prior to their geographic extinction from the Cape Flats. We generated complete mitochondrial and low-coverage nuclear genome-wide data for two Cape lions from the Cape Flats. We then compared the data from these Cape lions to historic lion populations that lived across Africa and in India. We investigate the population dynamics of Cape lion populations prior to their eradication using established genome metrics (heterozygosity and inbreeding) that have been used to study genomic signatures of population isolation and small population size and that have been linked to long-term population fitness and persistence (Lande, 1994; Lynch et al., 1995; Kirkpatrick and Jarne, 2000; Kyriazis et al., 2021). These metrics, combined with phylogenomic analysis of mitogenomes, allowed us to contextualize mitochondrial and nuclear genomic characteristics of Cape lions as part of expansive genomics patterns in lion populations across Africa and in India. In addition, we provide novel inferences and discussion based on our genomic results in light of historic events preceding the Cape lion’s demise.

## Methods and Materials

### Sample information

Bone and tooth samples were collected from two Cape lion specimens that are currently housed at the Field Museum of Natural History, Chicago (see Supplementary File 2 for more background on the Cape lions in this study). These include Cape lion 1 (Singer 1, JCK10711) and Cape lion 2 (Singer 2, JCK10712). For each lion, we collected 2 grams of bone powder by drilling into the skull/petrous bone, and 2 grams of dentine and pulp powder by drilling into a canine tooth. To maximize ancient DNA yield per lion, each sample was processed separately. However, shotgun sequencing data for both sample types were pooled for each lion during bioinformatic analyses.

### Molecular analysis

Ancient DNA extractions and genomic library preparation were conducted in the Malhi Ancient DNA Laboratory, at the Carl R. Woese Institute for Genomic Biology, University of Illinois at Urbana-Champaign (UIUC). This laboratory is dedicated exclusively to ancient DNA studies. DNA extraction and library preparation was done following the protocol described in (de Flamingh et al., 2022) and included negative controls to account for possible contamination by external DNA sources. In brief, whole genomic libraries were constructed using the NEBNext® Ultra II™ DNA Library Prep kit and NEBNext® Multiplex Oligos (Unique Dual Indexes) for Illumina®. The extracted DNA was pre-treated with USER (Uracil-Specific Excision Reagent) enzyme to remove cytosine to uracil nucleotide base changes that are common in ancient DNA (Hofreiter et al., 2001). All samples and negative libraries were pooled and shotgun sequenced to generate 150bp paired-end reads on an Illumina NovaSeq 6000 platform at the Roy J. Carver biotechnology center at the University of Illinois, Urbana-Champaign.

### Bioinformatic analysis

#### Mitochondrial DNA

Samples were de-multiplexed and the reads trimmed using the program *FASTP* v.0.19.6 (Chen et al., 2018) to have a minimum sequence length of 25bp. Reads were aligned to a reference lion mitogenome (PLE isolate with GenBank accession KP202262.1) reported by (Li et al., 2016) using *BOWTIE2* (Langmead and Salzberg, 2012). *BOWTIE2* alignments were converted into BAM format using *SAMTOOLS* view v. 1.1 (Li et al., 2009), and filtered to remove unmapped reads and reads with a quality score less than 30. Filtered BAM files were then sorted and indexed, with PCR duplicates marked and removed with the *PICARD TOOLKIT* v. 2.10.1 (Picard Toolkit 2019, Broad Institute). Mitochondrial read alignment datasets were filtered using *NUMT PARSER* (de Flamingh et al., 2022) to remove contamination from nuclear-mitochondrial elements. Consensus sequences were generated by retaining only regions that had at least 5X-fold read coverage, using the “Highest Quality” algorithm which considers relative residue quality to generate a majority consensus sequence in *GENEIOUS* (Kearse et al., 2012).

Cape lion mitogenomes were compared to mitogenome sequences from 118 lions from 13 African countries and from India, sampled between 1860 and the present day (Figure 1; Supplementary Table 1). Excluding three lions from contemporary populations, all other samples were reported to have been collected prior to 1950.

**Figure 1:**
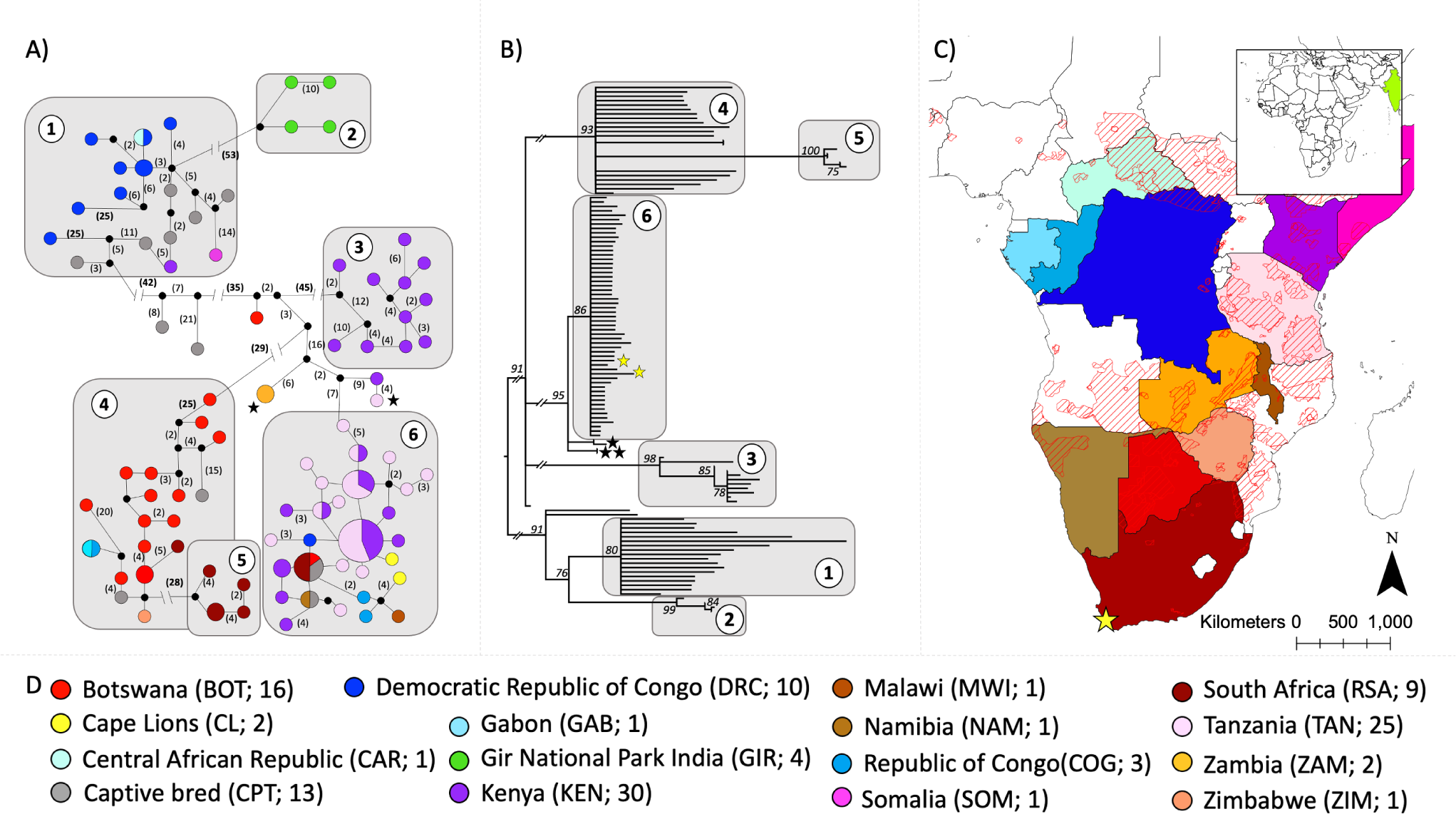
Phylogeographic analysis of two Cape lion and 114 African and 4 Indian lion mitogenomes. A median-joining analysis of the two complete Cape lion mitogenomes and 118 geographically referenced lion showed that 113 of the 120 lions broadly grouped into 6 clusters (panel A). A similar clustering pattern was observed in the maximum likelihood phylogeny, where 113 of 120 lions were partitioned into 6 corresponding clades (panel B). The geographic distribution of the samples is shown in panel C, and the location of historic Cape lion samples from the Cape Flats are indicated with a yellow star. Cape lion mitochondrial genomes grouped with a cluster (Cluster 6) that included lions from a wide geographic distribution, including southern and eastern Africa, but not with the cluster (Cluster 5) that is includes only lions from South Africa. Colors are consistent with geography across panels A-D (panel D).

Mitogenome sequences were aligned using the program *MUSCLE* (Edgar, 2004). The resulting alignment was then used to construct a median-joining network in the software *POPART* (Population Analysis with Reticulate Trees; Leigh and Bryant 2015), and to infer a maximum likelihood tree in *RAXML* v. 8 (Stamatakis, 2014) that was repeated 1000 times with 100 bootstrap iterations for each run, using two computational threads and a general time-reversible (GTR) model with optimization of substitution rates and a gamma model of rate of heterogeneity (GTRGAMMA; Stamatakis 2014).

#### Nuclear DNA analysis

The comparative nuclear DNA dataset comprised 53 lions from 14 African countries (Figure 2; Supplementary Table 2). To ensure similar sample representation from each country, we selected a subset of the lions from Curry et al. (2021) so that each country is represented by a maximum of 10 lions (Figure 2B). Unequal sample representation may bias estimates and interpretations of genetic variation (McVean 2009; Burgos-Paz et al. 2014).

**Figure 2:**
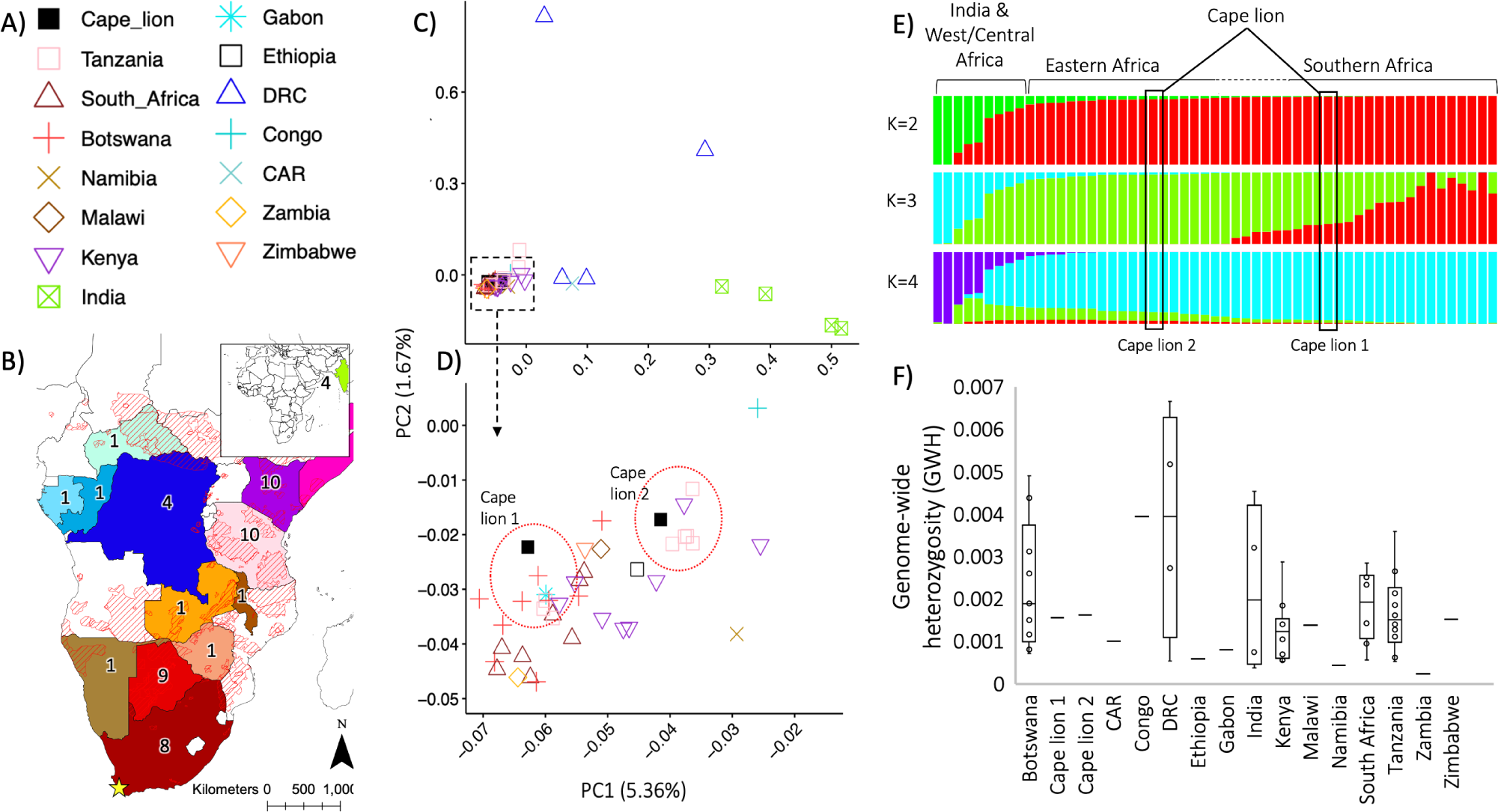
Analysis of genome-wide data from two Cape lions and 53 lions. These lions were from 14 countries (panel A) from across Africa (panel B) and from India (panel B - insert). In panel B, the numbers indicate the sample sizes for each country. In a principal component analysis (PCA) of genome-wide genetic variation (panel C), PC1 separated Asiatic lions from all other lions and positioned individuals from DRC and CAR as intermediate to Asiatic and other African lions, while PC2 separated two of four individuals from DRC from all other lions. Each principal component had a low contribution to the total variation; PC1 explains 5.36% of the variation, while other PCs explain <1.67% of the variation. When focusing in on the African lion cluster (panel D), PC1 followed a geographic cline with lions from southern Africa on the left transitioning to lions from eastern Africa on the right of PC1 (except for one individual from Namibia that was towards the right of PC1). Similarly, PC2 also followed a geographic cline with southern African lions occurring at lower values on the PC2 axis and transitioning into eastern African lions at higher values. Cape lion 1 and Cape lion 2 (panel C - red dashed circle) have distinct genomic profiles and respectively clustered with individuals from southern Africa, e.g., Botswana and towards the left in PC1 in the African lion cluster, and individuals from east Africa, e.g., Kenya and Tanzania towards the right of the PC1 axis in the African lion cluster. These distinct genomic profiles were also evident in the admixture analysis (panel E), where K=3 partitioned Cape lion 2 into a cluster that consisted of east African lions, while Cape lion 1 showed evidence of admixture between clusters that were from eastern and southern Africa. For both K = 2 and K = 3, India, West and Central Africa are partitioned into a single cluster separate from all other African lions. K = 4 did not show biogeographical partitioning beyond the separation of India, West and Central African lions from other African lions. The two Cape lions did not have depleted genome-wide heterozygosity or higher inbreeding coefficients compared to other historical populations (panel F). Colors in panels A-D consistently correspond to geographic location, while colors in panel E correspond to the probability of belonging to an admixture cluster (K).

Reads were aligned to the complete lion genome (v. PanLeo1.0, GenBank accession number GCA_008797005.1; Armstrong et al., 2020) using the mem module in *BWA* v. 0.7.15 (Li and Durbin, 2010). Alignments were converted to BAM format and filtered using the same criteria and software as for the mitochondrial analysis. We merged the BAM files from each sample type for each respective lion, and duplicate reads in merged files were marked and removed with the *PICARD TOOLKIT* v. 2.10.1. To verify that the genomic libraries had damage patterns that are consistent with ancient DNA, we quantified DNA damage in *MAPDAMAGE* v. 2.0.5 (Jónsson et al., 2013) using a fragment size of 70 bp. For each lion, we calculated the breadth of genome coverage (i.e., the percentage of the genome that has ≥1 X-fold read coverage) and the average depth of coverage (the average X-fold number of reads that mapped at any location across the genome). In addition, for each newly sequenced Cape lion, we calculated the depth of coverage distribution for each position across the genome using *SAMTOOLS* depth. Using a custom Python script (https://github.com/adeflamingh/de_Flamingh_etal_2023_Cape_lion), we processed the *SAMTOOLS* depth output file to calculate the mean depth of coverage within 250 Kbp-sized windows, every 100 Kbp (i.e., every 100 Kbp along the genome a new window of size 250 Kbp is calculated). The calculated averages for each lion were plotted across the genome using a custom R script (https://github.com/adeflamingh/de_Flamingh_etal_2023_Cape_lion) in R version 4.2.1 (Core Team R, 2013).

To investigate nuclear genome variation, we used a published phylogeographic analysis pipeline that was developed for low-coverage shotgun sequencing data (Yao et al., 2020). This pipeline uses genotype likelihoods rather than called genotypes to estimate single nucleotide polymorphisms (SNPs) in *ANGSD* (Korneliussen et al., 2014). The dataset only included SNPs that were present in at least half (n=28) of the individuals that had a significance (p-value) of ≤0.01, and with a minimum base quality of 30. We visualized the distribution of SNP variant sites along the chromosomes to ensure that the genetic variation observed for the dataset is representative of the complete lion genome. Similar to depth of coverage, we used a custom Python script (https://github.com/adeflamingh/de_Flamingh_etal_2023_Cape_lion) to count variant sites seen in the 55 individuals across 250 Kbp windows, again sampled every 100 Kbp. We confirmed that the variant sites retained were distributed along all autosomes and not localized to a limited region of the genome. Using a principal component analysis (PCA) in *PCANGSD* (Meisner & Albrechtsen, 2018), we investigated the genetic distances among individuals from different countries reported in published data (see Figure 2C for geographic distribution of samples per reported country), and we estimated admixture assuming 2, 3 and 4 clusters (K; Skotte et al. 2013). We repeated the PCA and admixture analyses for a subset of 46 lions that excluded lions from India and DRC to investigate genetic variation in Eastern and Southern African lion populations.

To investigate Cape lion population dynamics in the context of other lions from historic periods, and in consideration of historic events (e.g., the rapid eradication of wildlife from the Cape region), we calculated two genome metrics: genome-wide heterozygosity (GWH) and an inbreeding coefficient (ngsF). We estimated GWH in *ANGSD* as the proportion of heterozygous genotypes (analogous to theta-based estimates). Because the Cape lion DNA may contain changes that result from DNA degradation, e.g., deamination of cytosine bases (Rohland and Hofreiter, 2007), we considered only transversions when calculating GWH. Using the same genotype likelihood scores, we calculated per-individual inbreeding coefficients (F) in the program *NGSF* (Vieira et al., 2013). We used R to visualize the results of the PCA, admixture, GWH and ngsF analyses (source code available at https://github.com/adeflamingh/de_Flamingh_etal_2023_Cape_lion). Admixture proportions were iteratively sorted based on their probability of belonging to a cluster and their inbreeding coefficient was sorted based on their sample date, i.e., the most recently reported date when that sample was collected, prior to plotting.

We adapted the Rx method for genetic sex determination (Mittnik et al., 2016; de Flamingh et al., 2020) so that it can be used to estimate the genomic sex of lions. We tested the lion Rx method using published shotgun sequencing data from 5 male and 5 female historical lions (the adapted Rx code for lion sex determination is available at https://github.com/adeflamingh/de_Flamingh_etal_2023_Cape_lion). We then used this method to identify the genomic sex of both Cape lion specimens, and visualized genomic sex estimation results using R.

## Results

### Mitogenome analyses

We were able to reconstruct high coverage complete mitochondrial genomes for both Cape lions; we reconstructed 100% of the breadth of the mitogenome at 45.13 X-fold coverage of reads for Cape lion 1 and 31.98 X-fold coverage of reads for Cape lion 2. A median-joining analysis of the two complete Cape lion mitogenomes and 118 geographically referenced lion mitogenomes (Curry et al., 2021) showed that 113 of the 120 lions broadly grouped into 6 clusters (Figure 1A). A similar clustering pattern was observed in the maximum likelihood phylogeny, where 113 of 120 lions were partitioned into 6 corresponding clades (Figure 1B). The seven lions that did not group with clusters 1-6 in the median-joining network and maximum likelihood phylogeny included two captive lions, one lion from Botswana, two lions from Zambia (modern samples), one lion from Kenya and one lion from Tanzania (modern sample). The three lions that were from modern populations (Supplementary Table 1) did not group with any of the clusters in either the median-joining network or maximum likelihood phylogeny (Figure 1 – black filled stars). However, the two modern lions from Zambia did cluster together, and all three modern lion haplotypes were intermediately located between mtDNA clusters and grouped within a larger African lion clade on the maximum likelihood topology. Consistent with previous studies (Bertola et al., 2016; de Manuel et al., 2020; Curry et al., 2021) the mitochondrial DNA clustering pattern grouped lions from India with African lions from West Africa (cluster 1 and 2 in the median-joining network and maximum-likelihood phylogeny).

Clusters 1-5 showed geographic structuring where most individuals within a cluster were from a single country. The majority of individuals in Cluster 1 were from the Democratic Republic of the Congo (DRC) or were captive bred, and one lion each from Central African Republic (CAR), Kenya and Somalia. Cluster 1 therefore broadly includes lions from central to eastern Africa (see Figure1C). Cluster 2 included only individuals from the Gir National Park in India. Cluster 3 included only individuals from Kenya (eastern Africa). Most individuals in Cluster 4 were from Botswana, although one lion was from Zimbabwe and one haplotype was shared between two lions from Gabon and the Republic of Congo (ROC). Cluster 4 therefore broadly includes lions from central to southern Africa. Cluster 5 included only individuals from South Africa. Cluster 6 did not show phylogeographic structuring and comprised individuals from eight African countries, including Botswana, DRC, Kenya, Malawi, Namibia, ROC, South Africa, the two Cape Lions, Tanzania, and individuals bred in captivity (Figure 1). Although they were grouped together in Cluster 6, the two Cape lions had distinct mitogenome haplotypes. Cluster 2, which included only lions from Gir National Park in India, was separated by the most nucleotide base-pair differences (53 differences) in the median joining network (Figure 1A), supporting the position of Asiatic lions as an outgroup and as genetically distinct from African lions. The Cape lions did not group with Cluster 5 (which was geographically limited to South Africa) but rather grouped with Cluster 6 which does not follow phylogeographic structuring and includes individuals from across the African continent.

### Genome-wide analyses

To shed further light on the phylogeographic patterns in historical lion populations, we compared the nuclear genomes of the two Cape lions to genome-wide data from 53 lions across Africa and from India. We assembled 26.9% of the complete genome at an average depth of 0.33X-fold read coverage for Cape lion 1, and 21.3% of the genome at an average depth of 0.25X-fold read coverage for Cape lion 2 (Supplementary Table 3; Supplementary Figure 1). Reads were evenly distributed across the autosomes of Cape lion 1 and Cape lion 2, indicating that our dataset is representative of the complete historic lion genome (Supplementary Figure 1). Cape lion 1 had lower X-chromosome coverage which is consistent with the lion being male and carrying only a single X-chromosome (see the results for the Rx sex determination and Figure 3). In agreement, our Rx code for genomic sex estimation identified Cape lion 1 (Rx = 0.487) as male, while Cape lion 2 (Rx = 0.915) was identified as female. The adapted Rx code for lion genomic sex determination was able to accurately characterize the genomic sex of all 5 male and 5 female lions (Figure 3; Supplementary Table 5). Genotype likelihood analysis of the 55 lion genomes (Figure 2) resulted in the analysis of a total number of 2 364 730 445 sites, and after filtering, we retained 20 563 944 sites for subsequent analyses, e.g., PCA, GWH and inbreeding coefficient calculation. These filtered sites were consistently distributed across all autosomes (Supplementary Figure 2), while the X-chromosome showed lower coverage due to sex differences in the sampled individuals where genomically male individuals only carry one X-chromosome. The Cape lion DNA damage patterns showed nucleotide base-pair changes characteristic of ancient DNA, e.g., deamination of cytosine to uracil (Supplementary Figure 3; Hofreiter et al., 2001).

**Figure 3:**
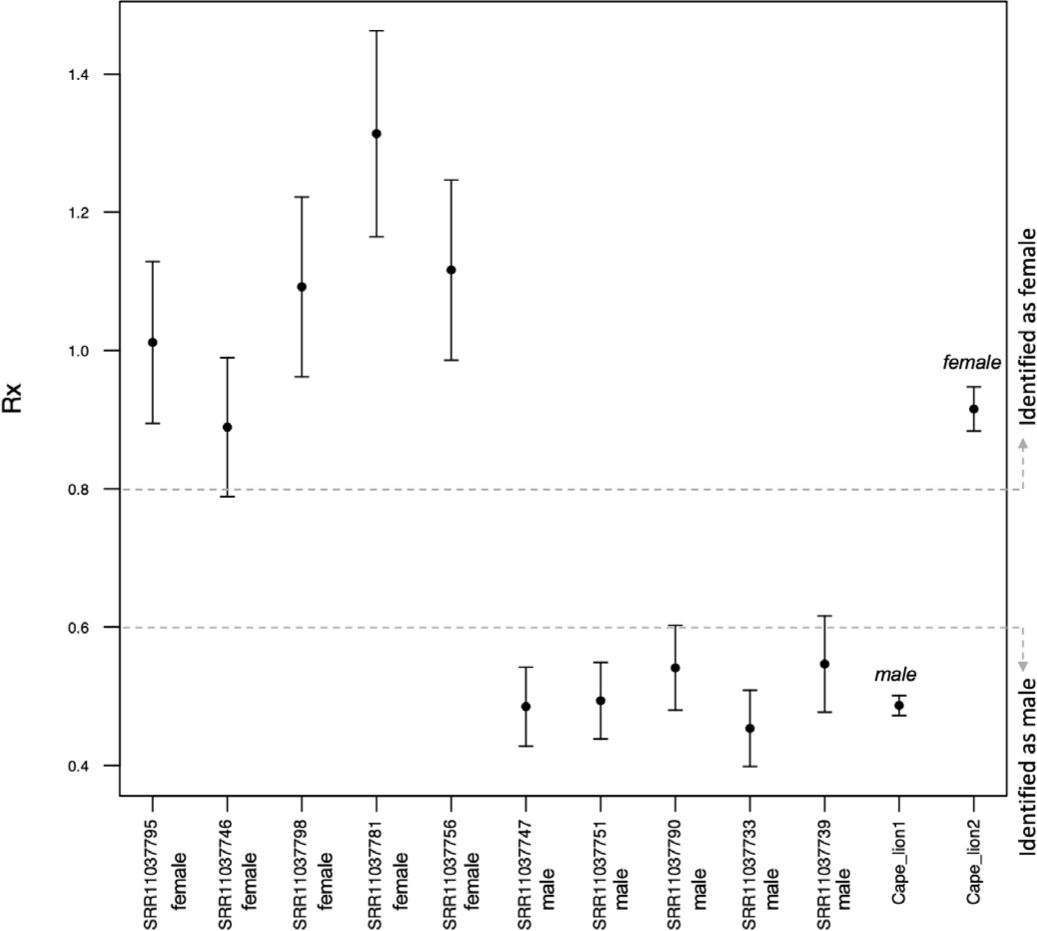
The adapted Rx code for lion genomic sex determination was able to accurately characterize the genomic sex of all 5 male and 5 female lions. The sequence read archive (SRA) accession numbers are indicated next to the reported genomic sex (Curry et al., 2019) on the x-axis. Cape lion 1 (Rx = 0.487) was identified as male, while Cape lion 2 (Rx = 0.915) was identified as female.

In a PCA of genome-wide genetic variation (Figure 2C), PC1 separated Asiatic lions from all other lions. Further, it and positioned individuals from DRC and CAR as intermediate to Asiatic and other African lions, while PC2 separated two of the four individuals from DRC from all other lions. Each principal component had a low contribution to the total variation; PC1 explains 5.36% of the variation, while other PCs explain ≤1.67% of the variation. When focusing on the African lion cluster (Figure 2D), PC1 followed a geographic cline with lions from southern Africa on the left, transitioning to lions from eastern Africa on the right of PC1 (except for one individual from Namibia that was towards the right of PC1). Similarly, PC2 also followed a geographic cline with southern African lions occurring at lower values on the PC2 axis and transitioning into eastern African lions at higher values. Considering only PC2, the Namibian lion (the outlier in PC1) clustered with other southern African lions. PC2 and PC3 separated two DRC and two Asiatic lions from other lions, but otherwise did not show geographic structuring within the African lion cluster. Our two Cape lions (Figure 2D right red dashed circle) have distinct genomic profiles: Cape Lion 1 clustered with individuals from southern Africa, e.g., Botswana and towards the left in PC1 in the African lion cluster, while Cape Lion 2 clustered with individuals from east Africa, e.g., Kenya and Tanzania towards the right of the PC1 axis in the African lion cluster. A repeated PCA and admixture analysis of a subset of 46 lions (Supplementary Figure 4), which excluded lions from India and DRC, showed similar structuring of lion genome-wide variation into southern to eastern geographic cline when considering PC1 and PC2. Cape lion 1 grouped with individuals from southern Africa, e.g., Malawi, Namibia, Zambia, while Cape lion 2 clustered with individuals from east Africa, e.g., Ethiopia, Kenya, Tanzania. These distinct genomic profiles were also evident in the admixture analysis (Figure 2E), where Cape lion K = 3 partitioned Cape lion 2 into a cluster that consisted of east African lions, while Cape lion 1 showed evidence of admixture between clusters that were from eastern and southern Africa while K=3 partitioned Cape lion 2 into a cluster that consisted of east African lions. This pattern is repeated when estimating admixture for the subset of 46 lions that excluded lions from DRC and India. For both K = 2 and K = 3 in the complete dataset (53 lions), India, West and Central Africa are partitioned into a single cluster separate from all other African lions (Figure 2E). K = 4 did not show biogeographical partitioning beyond the separation of India, West and Central African lions from other African lions.

The two Cape lions did not have depleted GWH or higher inbreeding coefficients compared to other historical populations (Figure 2F). Both Cape lions had similar GWH; Cape lion 2 (GWH = 0.00162), which clustered with southern African lions on PCA, had a slightly higher GWH than Cape lion 1 (GWH = 0.00156), which clustered with eastern African lions on PCA. Cape lion 1 and Cape lion 2 respectively had inbreeding coefficients (F) of 0.103 and 0.069 (Supplementary Table 4; Supplementary Figure 5), which are lower than the average F for historical lions from Africa (average F = 0.179), India (average F = 0.259), and modern lions (average F = 0.511).

## Discussion

Starting in the mid 1600’s concurrent with the introduction of sheep farming and the establishment of European settlements, Cape lions were eradicated from the Cape Flats region of South Africa (Bryden, 1889; Selous, 1908; Guggisberg, 1963). Simultaneous to the destruction of predators in the Cape and environs, there was also increased subsistence hunting of lion prey species, especially Springbok, that competed for grazing forage with sheep (Skead, Vol; 1 1980 & Vol. 2 1987 Thom, 1952). We show here that, before their eradication, lions from the Cape Flats had diverse mitochondrial haplotypes that clustered with other lions from southern and eastern Africa. In congruence, their nuclear genome-wide characteristics showed that these lions had different genomic profiles, despite being from the same geographic location. Previous research characterized patterns of nuclear genome variation across modern African lion populations and showed that there is a geographic “suture zone” that represents sympatric occurrence of lions with genome characteristics typical of eastern and of southern Africa (described as eastern and southern clades in Bertola et al., 2022). Nuclear genome analyses using microsatellite and SNP markers of modern lion populations indicated that this suture zone extends from northern Zambia down to central Mozambique. However, our analysis of genome-wide data of the two Cape lions suggests that the sympatric occurrence of eastern and southern nDNA clades may have historically extended as far south as the Cape Flats. Specifically, the genomic profile of Cape lion 2 matches historical lions from east Africa, while genomic analysis of Cape lion 1 shows admixture between lions from southern and eastern Africa. Population contiguity, connectivity and/or gene-flow, may therefore have historically extended across larger geographic scales than what has been reported for modern populations. Our findings thus do not support the characterization of Cape lions as a geographically and genetically distinct group, as was originally put forth by some European naturalists (Griffith, 1821; Smith, 1842; Mazak, 1964; Mazák, 1975), but are consistent with the molecular analyses of partial mitochondrial genomes reported by Barnett and colleagues (2006a, 2006b).

Of interest, Sparrman (1785) noted that resident breeding lions had been extirpated from much of the settled portions of the Western Cape by the 1780’s, especially in the areas closest to Cape Town and in environs where sheep were raised. According to Sparrman, however, lions continued to appear from time to time in the Western Cape ‘from the north’ (the Karoo), where they presumably followed an annual east-west migration (i.e., ‘Trekbokken’ in the Karoo), of Springbok, Cape Hartebeest, Eland, Black Wildebeest, Quagga and other prey species until both predator and prey were eventually eradicated (Skead, 1980). According to Gordon Cumming (1856), a local population of lions persisted in the Western Cape in the vicinity of the remaining intact migration of Trekbokken, in the Middleburg district of the Great Karoo between Graaff Rienet and Colesberg, up until the late 1840’s. The last known Cape Lion specimen record for the Western Cape, in 1848, is a lion cranium and mandible that is now at the Natural History Museum in London (catalog number BM.36.5.26.6). A historical annual migration of lion prey species, apparently similar in scale to the migration in the Serengeti ecosystem in northern Tanzania and southern Kenya (Schaller, 2009), would draw nomadic lions from geographically distant areas back into regions where resident breeding lions had been previously eradicated or were no longer present (Hanby and Bygott, 1979). Therefore, prey migration could potentially have created a functional landscape corridor that would have allowed for corresponding gene-flow and genomic connectivity between Eastern and Southern African lion populations. This is supported by studies on contemporary prides that show that seasonal lion space use is highly dependent on prey availability (Kittle et al., 2016). The disruption of such landscape corridors may have contributed to the increased genetic distinctiveness, lower gene-flow and lower genetic diversity in present-day regional lion populations (Bertola et al., 2022). In addition, the distinct phenotypic characteristics that some of these extinct Cape lions exhibited, e.g., dark and/or more elaborately developed manes (which are not necessarily correlated) similar to some current-day lion populations, may also have been in part a consequence of the variable environments in which they occurred, especially with regard to climate; temperature and humidity (Selous, 1908; Percival, 1927; Broschart and Gnoske, 2001; Gnoske et al., 2006). The presence of variable mane color enumerated as “…yellow, grey, and black.” and the movement of lions into unoccupied areas is also supported by historical accounts (Leslie, 1833).

Our genomic findings do not support some European naturalists’ historical type descriptions of Cape lions as comprising a distinct group that differed from other African lions (Griffith, 1821; Mazak, 1964; Mazák, 1975; Smith, 1842). Rather, our results are potentially consistent with alternative knowledge-systems e.g., traditional ecological knowledge and Indigenous perspectives (Agrawal, 1995; Huntington, 2000; Jessen et al., 2022) and how they can inform the evolutionary and ecological interpretation of species’ life histories. Indigenous artists made petroglyphs of lions with light or mixed color manes in the Western Cape Province where the Cape lions in this study are from (Figure 4A) and in the Eastern Cape Province which is thought to be part of the broader historical range of Cape lions (Figure 4 B,C; Skead, 1980). Art that was made by Indigenous peoples depicted lions and other recognizable species (e.g., eland and cattle) in renderings of pigments applied to stone surfaces. While the intended meaning and original purpose of these artistic renderings is subjective to the artist, the images of the lions that are portrayed depict the variation in mane color, size, and body color. These descriptions are also part of Indigenous oral histories that were transmitted to Europeans by the San and Khoisan Indigenous peoples who shared their considerable Indigenous historic knowledge with the first Dutch settler-pastoralists and other European colonists who followed (Burchell, 1824; Bennett, 1829). In addition to consulting written accounts of Indigenous knowledge or referencing Indigenous artworks, we encourage researchers to consult directly with living Indigenous communities and pursue research frameworks that are rooted in community engagement and community-based research (Claw et al., 2018; Schroeder et al., 2019; Tsosie and Claw, 2020).

**Figure 4:**
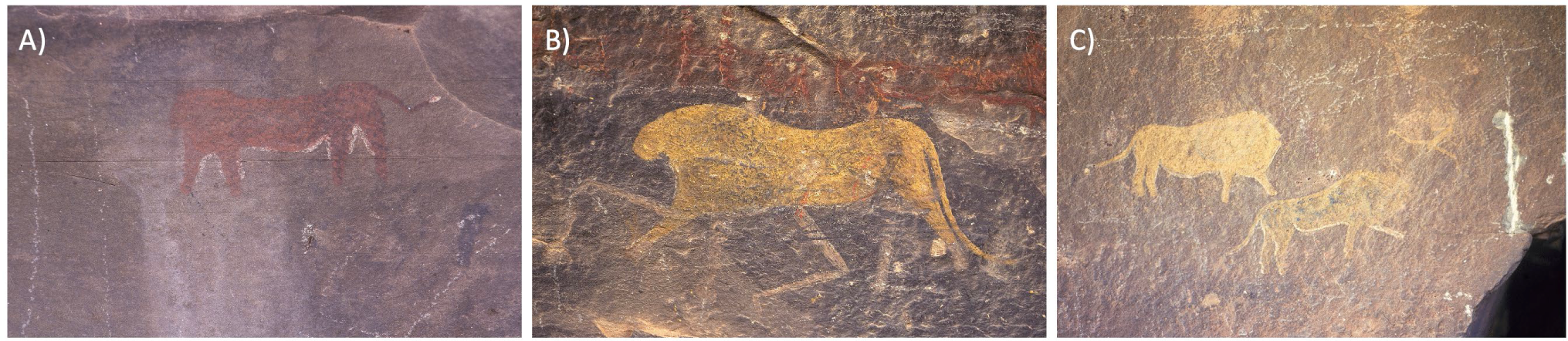
Petroglyphs created by Indigenous artists in what is today known as Murraysburg in the Western Cape Province (panel A), and the Wodehouse district in the Eastern Cape Province (panel B and C) of South Africa. Of note is the light coloration and lack of extensive black mane development. These images may represent the artist’s subjective interpretation rather than reflecting specific phenotypic variation, but together with other historical accounts point towards varied mane color in Cape lion populations. Usage permission of these images was granted through the South African Digital Rock Art Data Archive (SARADA), Rock Art Research Institute, University of Witwatersrand, South Africa; panel A (SARADA number JDC-RSA-PLX1-4) is the property of Janette Deacon, and panels B and C (SARADA image numbers RARI RSA DRI1 1 and RARI RSA BUF1 41) are the property of Jeremy Hollmann and RSA DRI1 site collection.

The only other genome-scale study of historic South African lions included two “Cape lions” (de Manuel et al., 2020) from geographically distinct parts of South Africa (Cape Town in the south, and King William’s Town which is ∼1000km east of Cape Town). However, genome-wide data support the placement of both these Cape lions within the genetic diversity found in South African lions. Since very few historical Cape lion genomes have been analyzed, it is feasible that the diversity in genomic profiles will increase as the genomes of other Cape lion specimens are sequenced.

The previous study (de Manuel et al., 2020) that included genome-wide analyses of two Cape lions investigated the long-term evolutionary history of extinct and living lions over a 30 000 year timeframe, whereas our study specifically contextualized Cape lion genome variation through comparison to historic lion populations from across Africa. We find that Cape lions had relatively intermediate genome-wide heterozygosity and low inbreeding coefficients compared to other historic lions, which suggests that the Cape lion population, and specifically the population in the Cape Flats, did not experience long-term isolation and small population size. In contemporary lion populations, habitat fragmentation and habitat destruction have drastically impacted population size and genomic diversity in both African (Bertola et al., 2016; Curry et al., 2021) and Asiatic (O’Brien et al., 1987) lions. Our analysis and comparisons of genome wide data from historical African, Asiatic, and modern lions also showed comparatively low heterozygosity and high inbreeding coefficients in modern lions. This is concerning because such decreases in genetic diversity have been associated with the expression of deleterious phenotypic characteristics in lions (e.g., abnormal sperm production Munson et al., 1996; Wildt et al., 1987). The genome-wide patterns of the two Cape lions in this study suggest that Cape lion populations in the Cape Flats went extinct so rapidly that genomic effects associated with long-term small population size and isolation were not visible. The rapid eradication of lion populations from the Cape Flats region during colonial settlement exemplifies the detrimental impact European colonization on African wildlife species and ecosystems (Ripple et al., 2015).

Understanding historical population dynamics and species’ genome characteristics preceding local population extinction can also inform contemporary conservation efforts for populations or species that may face similar threats. This can contribute to the success of wildlife conservation initiatives that aim to reintroduce populations to areas where a species previously ranged (He et al., 2016; Latch, 2020). Historical benchmarks may be especially beneficial for present-day lion population management, where there has been a call for the introduction of genomic diversity, e.g., translocation (Bertola et al., 2022).

In this study, we also provide an accessible and accurate method for determining the genomic sex of lions using next-generation shotgun sequencing data. The method is effective for estimating the genomic sex of even low-coverage historic DNA data and holds potential for anti-poaching strategies. Lion bones are a commodity that is part of the illegal wildlife trade (Williams et al., 2017), and the ability to characterize the genomic sex from lion bones could help anti-poaching strategies, e.g., by identifying whether male lions that usually roam alone across larger areas, or prides that include female lions and that live in less transitionary and more stable geographic areas, are being targeted by poachers. Our results therefore contribute towards understanding historic population dynamics of Cape lions to complement and inform current conservation initiatives that aim at protecting this vulnerable and declining species (Pečnerová et al., 2017; Curry et al., 2021; Bertola et al., 2022; IUCN Red List of Threatened Species, 2022).

## Acknowledgements

We want to highlight the important scientific contributions of the late Dr Ronald Singer, former Chair of the Anatomy Department at the University of Chicago, and a brilliant anatomist and paleoanthropologist. We thank Hazel and Eric Singer for access to and use of these historical specimens. We thank the UIUC High-Throughput Sequencing and Genotyping Unit. We thank Dr. Adam Ferguson and Dr. Tolulope Perrin-Stowe for their help in facilitating access to and collection of the historic DNA samples at the FMNH. For funding, we thank the USAID Wildlife TRAPS Project and the UIUC ACES Office of International Programs. AdeF was supported by the Program in Ecology, Evolution and Conservation Biology Research Award, UIUC, and by the Cooperative State Research, Education, and Extension Service, US Department of Agriculture, under project number ILLU 875–952. AGR-C was supported by NSF grant 1645087.

## Data Availability

## Bioinformatic code

https://github.com/adeflamingh/de_Flamingh_etal_2023_Cape_lion

## Genetic data

Raw sequence reads for the two Cape lions are deposited in the SRA (BioProject XXX). Unique mitogenome haplotype data are deposited to NCBI Nucleotide Database (XXXX).

## Author Contributions

AdF and RSM contributed to research design and execution, data generation, analysis, and interpretation of the genomics aspects of this study. ALR contributed towards the design, analysis, and interpretation of the genomics aspects of the study. TPG and JCK contributed to specimen acquisition, research design and data interpretation. VS and NY contributed to data interpretation. AGR-C, KW and JC contributed to the bioinformatic analyses. All authors contributed to the writing and editing of the manuscript.

