## Supplementary Files for "Genomic analysis supports Cape Lion population connectivity prior to colonial eradication and extinction"

##### *Supplementary File 1: Replicated descriptions and location of the Cape lion type specimen*

Consistent with Charles Hamilton Smith (Smith, 1842), the first detailed description of a wild South African lion from the Cape region was published by Andrew Smith in March 1834 (Smith, 1984). Andrew Smith had been director and superintendent of the South African Institute in Cape Town (1825-1837), the forerunner to the South African Museum established in 1855. The male lion described by Andrew Smith had a particularly well developed "black, brown and yellow mane" that covered its shoulders and abdomen and that varied only slightly from the wild-caught and captive raised individual described by Charles Hamilton Smith.

The specimen described by Andrew Smith was nearly identical in body dimensions to that of the first wild-collected adult Cape lion to be described, measured and weighed by a European, Jan van Riebeeck, on 16 June, 1656 (Thom, 1952), an average size for adult South African lions (Stevenson-Hamilton, 1947). These two early descriptions of wild-shot Cape Lion specimens are also consistent with a specimen of a late prime aged wild shot black maned male lion currently housed in the Natural History Museum in South Kensington London (BM.68.268), described and illustrated by R. I. Pocock (Pocock, 1931). It was thought to have originated from the then eastern boundary of the Eastern Cape, near the Orange River between 1830-1836 (Selous, 1908).

##### *Supplementary File 2: Background on the Cape lion specimens in this study*

The original label on the Singer Cape Lion skulls indicates they were at the "South African Museum", and came from a farm near the town of Phillipi in the Cape Flats, from the Dune-Strandveld/Sand Fynbos matrix, from the 1830's and would therefore be from the South African Institute of Andrew Smith <1825-1838>; the precursor of the South African Museum in Cape Town, established in 1855.

Their condition indicates that the skulls of the two Singer Cape lion specimens had been mounted inside of lion skins. This was a common practice during that period, and consistent with three other Maison Verreaux-family mounted lions that have been examined by three of us (TPG, JCKP and VS). In addition to other large mammals from the Colony, these Cape lion specimens were the same lion specimens prepared by 'Mr.' Jules Verreaux (Curator of the South African Institute 1829-1838) and brother Edouard, between 1834-1838. Following Smith's and Verreaux' departure (1837-1838) and permanent return to Europe (Summers, 1975), specimens from the collection were stored at Orphan House on Long Street, the first home of the South African College. Shortly thereafter, a naturalist visited the collection in October 1843 (Methuen, 1848), only five years after it closed to the public, and describes it as having fallen into a "lamentable state of wreck". These may represent the original male and female lion skins with crania mounted inside from A. Smith's South African Institute 1825-1838 (Layard, 1861). This is consistent with our genomic analysis in this study (Figure 3). E.L. Layard, the first curator of the newly formed South African Museum (1855-1872), began to remove or exchange the old and damaged materials that were from the South African Institute (Christiansen, 2008), including the mounted lion skins at some point, and eventually replaced those with newer material (Summers, 1975). These two lion crania appear to have been removed prior to the skins being discarded or disposed of by Layard sometime after 1862. There was no longer any record of these lion specimens being at the South African Museum by 1896, when W.L. Sclater (Director of the South African Museum 1896-1906) published 'The Mammals of South Africa (Sclater, 1900). According to the specimen tag, a Mrs. Johns had acquired the two lion crania, apparently long after they had been removed and disassociated from the South African Museum (between 1862 & 1896). She may have transmitted them to Dr. R. Singer, of the Department of Medicine at University of Cape Town, before Singer emigrated to Chicago and took a position at the University of Chicago (1962). Singer loaned the two skulls to one of us (JCKP) in 2003. Permission to access and study these historical Cape lion skulls were granted by Hazel and Eric Singer.

### Supplementary Figures

#### *Supplementary Figure 1*

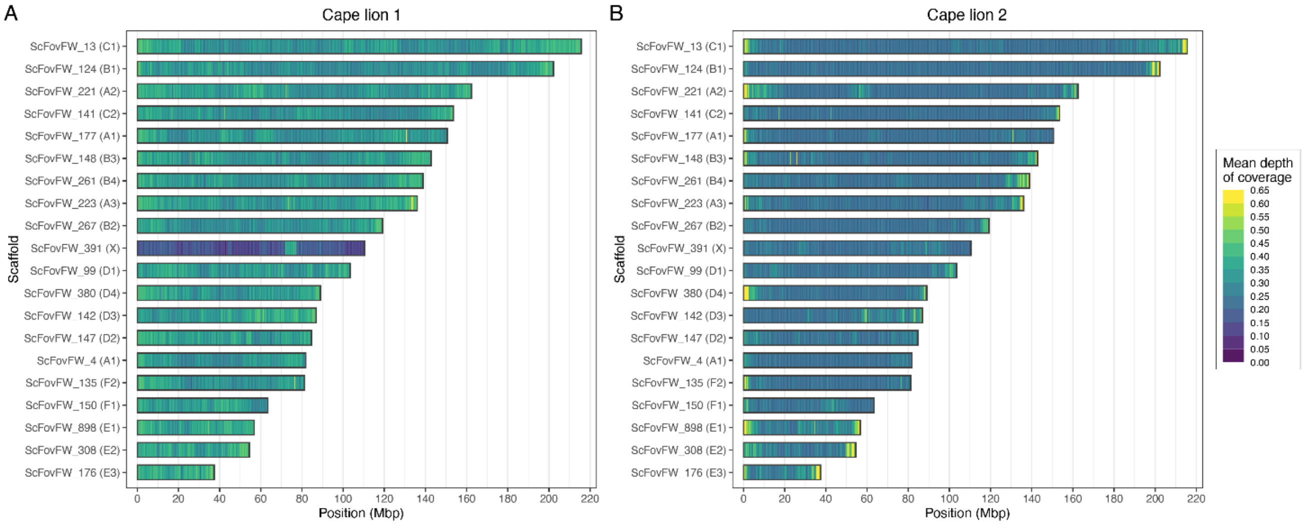


Read coverage distribution across the genome for Cape lion 1 (panel A) and Cape lion 2 (panel B). Each horizontal bar represents a chromosomal scaffold, and the x-axis shows the position on each chromosome. Colors represent the depth of coverage (X-fold) averaged across 250 Kbp windows along each chromosome. Cape lion 1, with a mean depth of coverage of ~0.33 X, shows reduced reads for the X-chromosome, which is consistent with its genomic sex estimation (see Rx sex determination and Figure 4) as being male and therefore carrying only a single copy of the X-chromosome. Cape lion 2 (estimated to be female), while having a lower average depth (~0.25 X), shows an even distribution of reads across all chromosomes.

##### *Supplementary Figure 2*

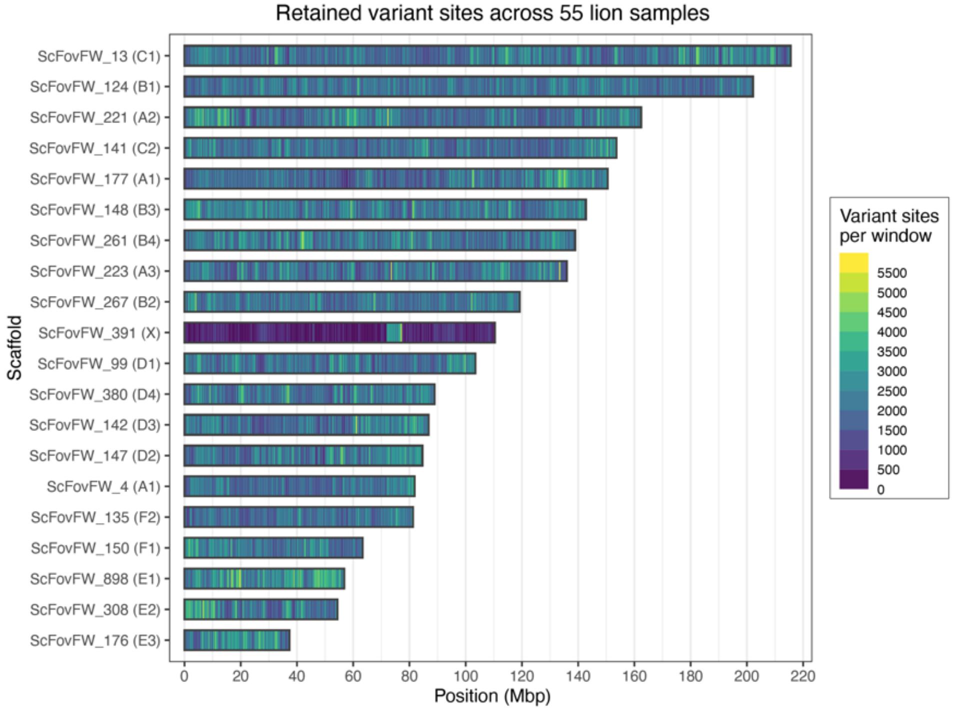


Distribution of SNP variant sites retained after filtering across the 55 lions included in this study (Supplementary Table 4, see main text for filtering criteria). Each horizontal bar denotes a chromosomal scaffold, with the x-axis showing the position along the chromosomes. Colored bars show the number of retained variant sites kept across a 250 Kbp window. On average, 2088 variant sites are seen in each window, except for the X chromosome (scaffold ScFovFW_391) in which only 678 sites are retained in each window (consistent with some individuals being male and therefore carrying only a single X-chromosome copy).

*Supplementary Figure 3*

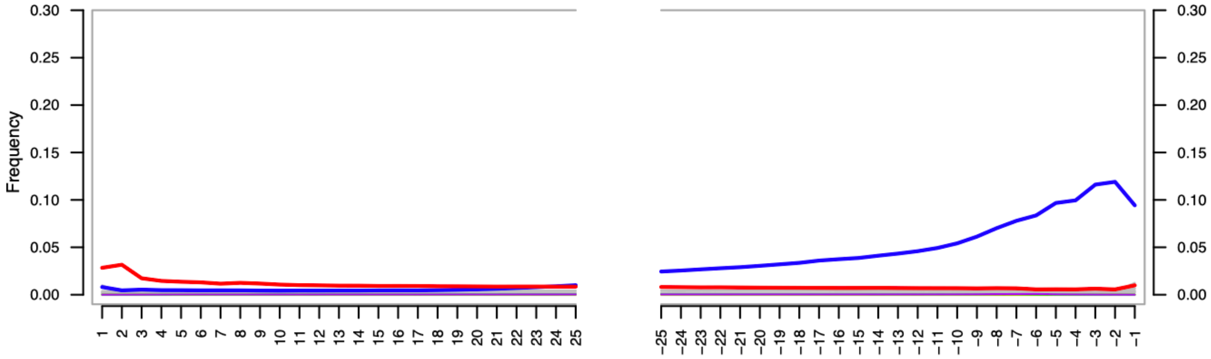


The Cape lion DNA showed damage patterns characteristic of ancient DNA (e.g., deamination of cytosine to uracil, (Hofreiter et al., 2001). A fragment misincorporation plot is shown for Cape lion 1 where red indicates C to T transitions, blue indicates G to A transitions. The y-axis denotes frequency of nucleotide change from the reference sequence, and x-axis denotes position along the DNA fragment with the left side of the x-axis showing nucleotides from the 5’ end of the read going into the read from left to right, and the right side of the x-axis showing nucleotides from the 3’ end of the read going into the read from right to left. This figure was produced using the program mapDamage2 (Jónsson et al., 2013). The damage patterns were similar for Cape lion specimen Singer No 2 (not shown).

***Supplementary Figure 4***

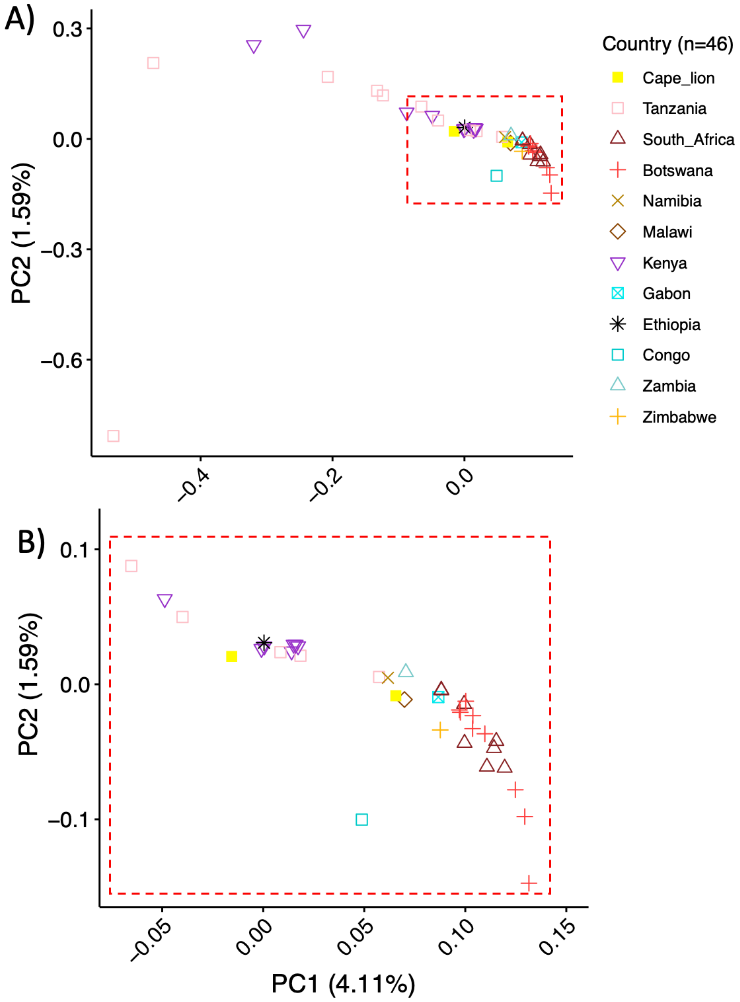


A PCA of genetic variation of 46 lions (A), which excluded lions from India and DRC, also showed the partitioning of lions into a geographic cline. An enlargement of the main cluster containing southern African lions (B) showed that the Cape lions had unique genomic profiles, with Cape lion 2 clustering with individuals from east Africa and Cape lion 1 clustering with individuals from southern Africa.

#### *Supplementary Figure 5*

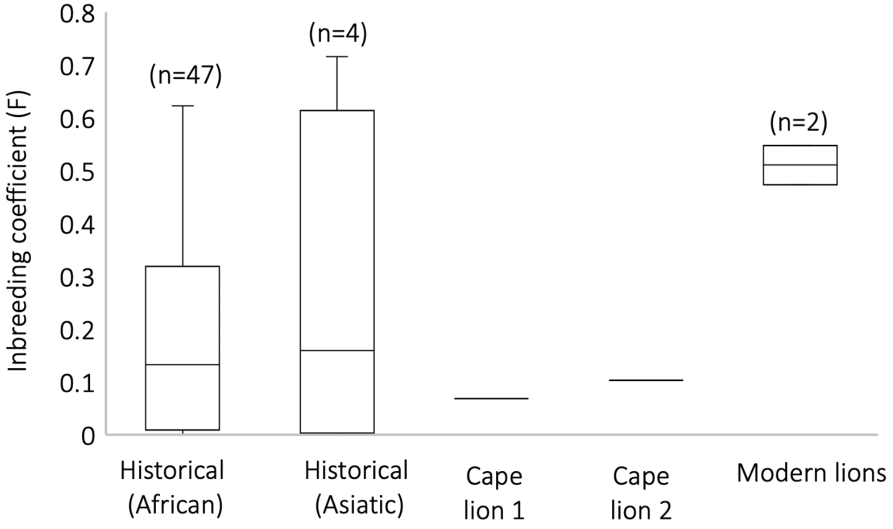


Using the same genotype likelihood scores for 55 lions from across Africa, we calculated the per-individual inbreeding coefficients (F) in the program ngsF (Vieira et al., 2013). Cape lion 1 and Cape lion 2 respectively had inbreeding coefficients (F) of 0.103 and 0.069, which are lower than the average F for historical lions from Africa (average F = 0.179), India (average F = 0.259), and modern lions (average F = 0.511), indicating that the Cape lion population did not show evidence of inbreeding compared to other historical or modern lion populations.

### Supplementary Tables

#### *Supplementary Table 1*

The mitochondrial DNA dataset included complete mitogenomes for 120 lions, 118 of which were previously published (Curry et. al., 2020) and 2 historic Cape lion specimens that were sequenced in this study.

| Sample number | Published_name | Museum ID | Origin | Collection year |
|---|---|---|---|---|
| 1 | H_BOT_001 | AMNH_11 9594 | BOT | 1930 |
| 2 | H_BOT_002 | AMNH_11 9595 | BOT | 1930 |
| 3 | H_BOT_069 | AMNH_83617 | BOT | 1930 |
| 4 | H_BOT_070 | AMNH_83618 | BOT | 1930 |
| 5 | H_BOT_071 | AMNH_83619 | BOT | 1930 |
| 6 | H_BOT_072 | AMNH_83620 | BOT | 1930 |
| 7 | H_BOT_073 | AMNH_83621 | BOT | 1930 |
| 8 | H_BOT_074 | AMNH_83622 | BOT | 1930 |
| 9 | H_BOT_075 | AMNH_83623 | BOT | 1930 |
| 10 | H_BOT_076 | AMNH_83624 | BOT | 1930 |
| 11 | H_BOT_077 | AMNH_83625 | BOT | 1930 |
| 12 | H_BOT_127 | FMNH_35739 | BOT | 1930 |
| 13 | H_BOT_129 | FMNH_3574 1 | BOT | 1930 |
| 14 | H_BOT_131 | FMNH_35 74 3 | BOT | 1930 |
| 15 | H_BOT_133 | FMNH_41405 | BOT | 1930 |
| 16 | H_BOT_137 | FMNH_8992 6 | BOT | 1935 |
| 17 | H_CAR_067 | AMNH_83410 | CAR | 1924 |
| 18 | H_COG_003 | AMNH_119870 | COG | 1949 |
| 19 | H_COG_012 | AMNH_172 74 | COG | <1920 |
| 20 | H_COG_013 | AMNH_172 75 | COG | <1920 |
| 21 | H_CPT_006 | A M NH_1 3998 | CAP | 1898 |
| 22 | H_CPT_007 | AMNH_1402 7 | CAP | 1895 |
| 23 | H_CPT_008 | AMNH_14 02 8 | CAP | 1895 |
| 24 | H_CPT_015 | AMNH_24 249 | CAP | 1905 |
| 25 | H_CPT_052 | AMNH_6260 | CAP | 1893 |
| 26 | H_CPT_053 | AMNH_6282 | CAP | 1889 |
| 27 | H_CPT_055 | AMNH_65 | CAP | 1860 |
| 28 | H_CPT_056 | AMNH_70171 | CAP | 1924 |
| 29 | H_CPT_097 | CM_1461 | CAP | No date |
| 30 | H_CPT_098 | CM_1564 | CAP | 1908 |
| 31 | H_CPT_099 | CM_1565 | CAP | 1908 |
| 32 | H_CPT_100 | CM_1825 | CAP | No date |
| 33 | H_CPT_158 | YPM_6943 | CAP | No date |
| 34 | H_DRC_031 | AMNH_52 07 0 | DRC | 1911 |
| 35 | H_DRC_033 | AMNH_S2072 | DRC | 1911 |
| 36 | H_DRC_034 | A M NH_ 52073 | DRC | 1911 |
| 37 | H_DRC_035 | AMNH_5 207 4 | DRC | 1911 |
| 38 | H_DRC_036 | AMNH_52 075 | DRC | 1911 |
| 39 | H_DRC_038 | AMNH_52077 | DRC | 1912 |
| 40 | H_DRC_039 | AMNH_52078 | DRC | 1912 |
| 41 | H_DRC_040 | AMNH_52079 | DRC | 1912 |
| 42 | H_DRC_042 | AMNH_52081 | DRC | 1912 |
| 43 | H_DRC_043 | AMNH_52082 | DRC | 1912 |
| 44 | H_GAB_004 | AMNH_119871 | GAB | 1949 |
| 45 | H_GIR_050 | AMNH_54995 | GIR | 1929 |
| 46 | H_GIR_051 | AMNH_54996 | GIR | 1929 |

| | | | | |
|---|---|---|---|---|
| 47 | H_GIR_054 | AMNH_63955 | GIR | 1906 |
| 48 | H_GIR_120 | FMNH_31121 | GIR | 1929 |
| 49 | H_KEN_016 | A M NH_277 69 | KEN | 1906 |
| 50 | H_KEN_019 | AMNH_3 024 1 | KEN | 1912 |
| 51 | H_KEN_020 | AMNH_30242 | KEN | 1912 |
| 52 | H_KEN_021 | AMNH_302 43 | KEN | 1912 |
| 53 | H_KEN_022 | AMNH_30244 | KEN | 1912 |
| 54 | H_KEN_023 | A M NH_30245 | KEN | 1912 |
| 55 | H_KEN_024 | AMNH_3024 6 | KEN | 1912 |
| 56 | H_KEN_025 | AMNH_302 47 | KEN | 1912 |
| 57 | H_KEN_026 | AMNH_3024 8 | KEN | 1912 |
| 58 | H_KEN_028 | AMNH_36420 | KEN | 1912 |
| 59 | H_KEN_029 | AMNH_3642 1 | KEN | 1912 |
| 60 | H_KEN_044 | AMNH_54370 | KEN | 1912 |
| 61 | H_KEN_045 | AMNH_54371 | KEN | 1912 |
| 62 | H_KEN_046 | AMNH_54372 | KEN | 1912 |
| 63 | H_KEN_057 | AMNH_70347 | KEN | 1922 |
| 64 | H_KEN_091 | AMNH_88632 | KEN | 1912 |
| 65 | H_KEN_092 | AMNH_88633 | KEN | 1912 |
| 66 | H_KEN_094 | AMNH_88635 | KEN | 1912 |
| 67 | H_KEN_095 | AMNH_88636 | KEN | 1912 |
| 68 | H_KEN_096 | AMNH_88637 | KEN | 1912 |
| 69 | H_KEN_114 | FMNH_20756 | KEN | 1905 |
| 70 | H_KEN_115 | FMNH_20757 | KEN | 1905 |
| 71 | H_KEN_116 | FMNH_20758 | KEN | 1905 |
| 72 | H_KEN_117 | FMNH_20 760 | KEN | 1905 |
| 73 | H_KEN_118 | FMNH_20762 | KEN | 1905 |
| 74 | H_KEN_142 | LACM_512 95 | KEN | 1922 |
| 75 | H_KEN_154 | YPM_3189 | KEN | 1931 |
| 76 | H_KEN_155 | YPM_3217 | KEN | 1931 |
| 77 | H_KEN_156 | YPM_3218 | KEN | 1931 |
| 78 | H_KEN_157 | YPM_5251 | KEN | 1922 |
| 79 | H_MWI_011 | AMNH_161732 | MWI | 1946 |
| 80 | H_NAM_014 | AMNH_191 81 | NAM | <1913 |
| 81 | H_RSA_059 | AMNH_81836 | RSA | 1930 |
| 82 | H_RSA_060 | AMNH_81837 | RSA | 1930 |
| 83 | H_RSA_061 | AMNH_81839 | RSA | 1930 |
| 84 | H_RSA_062 | AMNH_81840 | RSA | 1930 |
| 85 | H_RSA_063 | AMNH_81841 | RSA | 1930 |
| 86 | H_RSA_064 | AMNH_81842 | RSA | 1930 |
| 87 | H_RSA_065 | AMNH_81843 | RSA | 1930 |
| 88 | H_RSA_066 | AMNH_81844 | RSA | 1930 |
| 89 | H_RSA_132 | FMNH_381 34 | RSA | 1928 |
| 90 | H_SOM_113 | FMNH_1443 | SOM | 1896 |
| 91 | H_TAN_079 | AMNH_85140 | TAN | 1928 |
| 92 | H_TAN_080 | AMNH_85141-L | TAN | 1928 |
| 93 | H_TAN_081 | AMNH_85141-N | TAN | 1928 |
| 94 | H_TAN_083 | AMNH_85142-N | TAN | 1928 |
| 95 | H_TAN_084 | AMNH_85143 | TAN | 1928 |
| 96 | H_TAN_085 | AMNH_85144 | TAN | 1928 |
| 97 | H_TAN_086 | AMNH_85145 | TAN | 1928 |
| 98 | H_TAN_087 | AMNH_85146 | TAN | 1928 |
| 99 | H_TAN_088 | AMNH_85147 | TAN | 1928 |
| 100 | H_TAN_089 | AMNH_85148 | TAN | 1928 |
| 101 | H_TAN_090 | AMNH_85149 | TAN | 1928 |
| 102 | H_TAN_105 | CM_5897 | TAN | 1927 |

| | | | | |
|---|---|---|---|---|
| 103 | H_TAN_106 | CM_5898 | TAN | 1927 |
| 104 | H_TAN_107 | CM_5899 | TAN | 1927 |
| 105 | H_TAN_108 | FMNH _127836 | TAN | 1928 |
| 106 | H_TAN_111 | FMNH_127839 | TAN | 1928 |
| 107 | H_TAN_123 | FMNH _35131 | TAN | 1930 |
| 108 | H_TAN_124 | FMNH_3513 2 | TAN | 1930 |
| 109 | H_TAN_125 | FMNH_3513 3 | TAN | 1930 |
| 110 | H_TAN_126 | FMNH_35134 | TAN | 1930 |
| 111 | H_TAN_140 | KU_1 05216 | TAN | 1920 |
| 112 | H_TAN_141 | KU_1 05217 | TAN | 1920 |
| 113 | H_TAN_145 | M VZ_9 68 04 | KEN | 1929 |
| 114 | H_TAN_153 | YPM_2057 | TAN | 1928 |
| 115 | H_ZIM _010 | AMNH_16101 1 | ZIM | 1946 |
| 116 | TAN_TLS | Modern | TAN | Modern |
| 117 | ZAM_T256 | Modern | ZAM | Modern |
| 118 | ZAM_T429 | Modern | ZAM | Modern |
| 119 | Cape lion 1 | JCK10711 | RSA | <1890 |
| 120 | Cape lion 2 | JCK10712 | RSA | <1890 |

#### *Supplementary Table 2*

The nuclear DNA dataset included genomic data from 53 previously published lion genomes (Curry et. al., 2020) from across the African continent, and two newly sequenced Cape lion genomes from the "Cape Flats" region of South Africa.

| Sample number | Individual Accession/ID | Location | Published ID | Reported sex | Collection date |
|---|---|---|---|---|---|
| 1 | SRR11037774 | DRC | H_DRC_040 | male | 1912 |
| 2 | SRR11037711 | Malawi | H_MWI_011 | female | 1946 |
| 3 | SRR11037783 | Congo | H_COG_013 | not collected | <1920 |
| 4 | SRR11037746 | India | H_GIR_051 | female | 1929 |
| 5 | SRR11037798 | Botswana | H_BOT_002 | female | 1930 |
| 6 | SRR11037747 | India | H_GIR_050 | male | 1929 |
| 7 | SRR11037779 | DRC | H_DRC_034 | female | 1911 |
| 8 | SRR11037781 | DRC | H_DRC_031 | female | 1911 |
| 9 | SRR11037756 | Tanzania | H_TAN_124 | female | 1930 |
| 10 | SRR11037788 | Botswana | H_BOT_133 | male | 1930 |
| 11 | SRR11037701 | South Africa | H_RSA_066 | not collected | 1930 |
| 12 | SRR11037744 | India | H_GIR_120 | male | 1929 |
| 13 | SRR11037750 | Botswana | H_BOT_070 | female | 1930 |
| 14 | SRR11037787 | Botswana | H_BOT_137 | not collected | 1935 |
| 15 | JCK10712 | Cape lion 2 | Cape_Lion2 | female | <1890 |
| 16 | SRR11037745 | India | H_GIR_054 | not collected | 1906 |
| 17 | SRR11037771 | Tanzania | H_TAN_083 | not collected | 1928 |
| 18 | SRR11037773 | DRC | H_DRC_042 | female | 1912 |
| 19 | SRR11037752 | Tanzania | H_TAN_141 | female | 1920 |
| 20 | SRR11037700 | South Africa | H_RSA_132 | male | 1928 |
| 21 | SRR11037706 | Botswana | H_BOT_074 | not collected | 1930 |
| 22 | SRR11037795 | Zimbabwe | H_ZIM_010 | female | 1946 |
| 23 | SRR11037761 | Botswana | H_BOT_076 | not collected | 1930 |
| 24 | SRR11037712 | Kenya | H_KEN_157 | not collected | 1922 |
| 25 | SRR11037715 | Kenya | H_KEN_154 | female | 1931 |
| 26 | SRR11037770 | Tanzania | H_TAN_084 | male | 1928 |
| 27 | SRR11037748 | Gabon | H_GAB_004 | not collected | 1949 |
| 28 | SRR11037790 | Botswana | H_BOT_129 | male | 1930 |
| 29 | SRR11037733 | Kenya | H_KEN_028 | male | 1912 |
| 30 | SRR11037743 | Kenya | H_KEN_016 | not collected | 1906 |
| 31 | JCK10711 | Cape lion 1 | Cape_Lion1 | male | <1890 |
| 32 | SRR11037794 | Tanzania | M_TAN_135 | male | modern |
| 33 | SRR11037769 | Tanzania | H_TAN_085 | male | 1928 |
| 34 | SRR11037749 | Ethiopia | H_ETH_104 | male | 1912 |
| 35 | SRR11037705 | South Africa | H_RSA_062 | female | 1930 |
| 36 | SRR11037696 | Tanzania | H_TAN_081 | not collected | 1928 |
| 37 | SRR11037713 | Kenya | H_KEN_156 | male | 1931 |
| 38 | SRR11037724 | Kenya | H_KEN_094 | not collected | 1912 |
| 39 | SRR11037786 | CAR | H_CAR_067 | not collected | 1924 |
| 40 | SRR11037709 | South Africa | H_RSA_059 | not collected | 1930 |
| 41 | SRR11037710 | Namibia | H_NAM_014 | female | <1913 |
| 42 | SRR11037708 | South Africa | H_RSA_060 | male | 1930 |
| 43 | SRR11037714 | Kenya | H_KEN_155 | not collected | 1931 |
| 44 | SRR11037703 | South Africa | H_RSA_064 | not collected | 1930 |
| 45 | SRR11037751 | Tanzania | H_TAN_145 | male | 1929 |
| 46 | SRR11037730 | Kenya | H_KEN_045 | not collected | 1912 |
| 47 | SRR11037717 | Botswana | H_BOT_073 | not collected | 1930 |
| 48 | SRR11037707 | South Africa | H_RSA_061 | female | 1930 |

| | | | | | |
|---|---|---|---|---|---|
| 49 | SRR11037726 | Kenya | H_KEN_091 | not collected | 1912 |
| 50 | SRR11037697 | Tanzania | H_TAN_080 | female | 1928 |
| 51 | SRR11037767 | Tanzania | H_TAN_087 | not collected | 1928 |
| 52 | SRR11037731 | Kenya | H_KEN_044 | not collected | 1912 |
| 53 | SRR11037739 | Botswana | H_BOT_071 | male | 1930 |
| 54 | SRR11037702 | South Africa | H_RSA_065 | not collected | 1930 |
| 55 | SRR11037793 | Zambia | M_ZAM_T256 | male | modern |

##### *Supplementary Table 3*

Mitochondrial and nuclear genome alignment statistics for Cape lion 1 and Cape lion 2. We calculated the breadth of genome coverage (the percentage of the lion genome that have >1X-fold read coverage) and the average depth of coverage (the average X-fold number of reads that mapped at any location across the genome).

| Mitogenome | Breadth of coverage (%) | Depth of coverage (X) | Reads (n) |
|---|---|---|---|
| Cape lion 1 | 100 | 45.13 | 11879 |
| Cape lion 2 | 100 | 31.98 | 8070 |

| Complete genomes | Breadth of coverage (%) | Depth of coverage (X) | Reads (n) |
|---|---|---|---|
| Cape lion 1 | 26.9368 | 0.329398 | 12946950 |
| Cape lion 2 | 21.2959 | 0.253813 | 9089619 |

#### *Supplementary Table 4*

Using the same genotype likelihood scores for 55 lions from across Africa, we calculated the per-individual inbreeding coefficients (F) in the program ngsF (Vieira et al., 2013). Cape lion 1 and Cape lion 2 respectively had inbreeding coefficients (F) of 0.103 and 0.069, which are lower than the average F for historical lions from Africa (average F = 0.179), India (average F = 0.259), and modern lions (average F = 0.511), indicating that the Cape lion population did not show evidence of inbreeding compared to other historical or modern lion populations.

| Individual ID | pop | Date | Inbreeding coefficient (F) |
|---|---|---|---|
| Cape lion 1 | Cape lion | Cape lion | 0.103247 |
| Cape lion 2 | Cape lion | Cape lion | 0.068700 |
| SRR11037696 | Tanzania | Historical (Africa) | 0.055459 |
| SRR11037697 | Tanzania | Historical (Africa) | 0.169743 |
| SRR11037700 | South Africa | Historical (Africa) | 0.025855 |
| SRR11037701 | South Africa | Historical (Africa) | 0.043603 |
| SRR11037702 | South Africa | Historical (Africa) | 0.245596 |
| SRR11037703 | South Africa | Historical (Africa) | 0.099417 |
| SRR11037705 | South Africa | Historical (Africa) | 0.009364 |
| SRR11037706 | Botswana | Historical (Africa) | 0.252307 |
| SRR11037707 | South Africa | Historical (Africa) | 0.420926 |
| SRR11037708 | South Africa | Historical (Africa) | 0.003662 |
| SRR11037709 | South Africa | Historical (Africa) | 0.079309 |
| SRR11037710 | Namibia | Historical (Africa) | 0.622948 |
| SRR11037711 | Malawi | Historical (Africa) | 0.319599 |
| SRR11037712 | Kenya | Historical (Africa) | 0.479774 |
| SRR11037713 | Kenya | Historical (Africa) | 0.461361 |
| SRR11037714 | Kenya | Historical (Africa) | 0.391159 |
| SRR11037715 | Kenya | Historical (Africa) | 0.346796 |
| SRR11037717 | Botswana | Historical (Africa) | 0.315009 |
| SRR11037724 | Kenya | Historical (Africa) | 0.007387 |
| SRR11037726 | Kenya | Historical (Africa) | 0.036166 |
| SRR11037730 | Kenya | Historical (Africa) | 0.467770 |
| SRR11037731 | Kenya | Historical (Africa) | 0.438852 |
| SRR11037733 | Kenya | Historical (Africa) | 0.206395 |
| SRR11037739 | Botswana | Historical (Africa) | 0.004326 |
| SRR11037743 | Kenya | Historical (Africa) | 0.004222 |
| SRR11037748 | Gabon | Historical (Africa) | 0.296141 |
| SRR11037749 | Ethiopia | Historical (Africa) | 0.087403 |
| SRR11037750 | Botswana | Historical (Africa) | 0.222408 |
| SRR11037751 | Tanzania | Historical (Africa) | 0.510554 |
| SRR11037752 | Tanzania | Historical (Africa) | 0.013486 |
| SRR11037756 | Tanzania | Historical (Africa) | 0.000000 |
| SRR11037761 | Botswana | Historical (Africa) | 0.431949 |
| SRR11037767 | Tanzania | Historical (Africa) | 0.037937 |
| SRR11037769 | Tanzania | Historical (Africa) | 0.137620 |
| SRR11037770 | Tanzania | Historical (Africa) | 0.138856 |
| SRR11037771 | Tanzania | Historical (Africa) | 0.168535 |
| SRR11037773 | DRC | Historical (Africa) | 0.023727 |
| SRR11037774 | DRC | Historical (Africa) | 0.403594 |
| SRR11037779 | DRC | Historical (Africa) | 0.000016 |
| SRR11037781 | DRC | Historical (Africa) | 0.000000 |
| SRR11037783 | Congo | Historical (Africa) | 0.007191 |
| SRR11037786 | CAR | Historical (Africa) | 0.233648 |

| | | | |
|---|---|---|---|
| SRR11037787 | Botswana | Historical (Africa) | 0.001957 |
| SRR11037788 | Botswana | Historical (Africa) | 0.009333 |
| SRR11037790 | Botswana | Historical (Africa) | 0.002772 |
| SRR11037795 | Zimbabwe | Historical (Africa) | 0.133266 |
| SRR11037798 | Botswana | Historical (Africa) | 0.025704 |
| SRR11037744 | India | Historical (Asiatic) | 0.000646 |
| SRR11037745 | India | Historical (Asiatic) | 0.010899 |
| SRR11037746 | India | Historical (Asiatic) | 0.715912 |
| SRR11037747 | India | Historical (Asiatic) | 0.309626 |
| SRR11037793 | Zambia | Modern | 0.548460 |
| SRR11037794 | Tanzania | Modern | 0.474119 |

#### *Supplementary Table 5*

The adapted Rx code for lion genomic sex determination was able to accurately characterize the genomic sex of all 5 male and 5 female lions. The sequence read archive (SRA) accession numbers are indicated next to the reported genomic sex (Curry et al., 2019) on the x-axis. Cape lion 1 (Rx = 0.487) was identified as male, while Cape lion 2 (Rx = 0.915) was identified as female.

| Name | Rx | Upper CI | lower CI | Sex det | Historically reported sex |
|---|---|---|---|---|---|
| SRR11037795 | 1.011582 | 0.894651 | 1.128512 | Sex assignment:The sample is consistent with XX but not XY | female |
| SRR11037746 | 0.8890354 | 0.7886902 | 0.9893805 | Sex assignment:The sample is consistent with XX but not XY | female |
| SRR11037798 | 1.091958 | 0.9618044 | 1.222112 | Female | female |
| SRR11037781 | 1.313582 | 1.164529 | 1.462635 | Female | female |
| SRR11037756 | 1.116277 | 0.9859642 | 1.24659 | Female | female |
| SRR11037747 | 0.4850119 | 0.4277121 | 0.5423118 | Male | male |
| SRR11037751 | 0.4936757 | 0.4383419 | 0.5490094 | Male | male |
| SRR11037790 | 0.5410463 | 0.4797153 | 0.6023772 | Sex assignment:The sample is consistent with XY but not XX | male |
| SRR11037733 | 0.4534757 | 0.3984276 | 0.5085238 | Male | male |
| SRR11037739 | 0.5466134 | 0.4771274 | 0.6160995 | Sex assignment:The sample is consistent with XY but not XX | male |
| Cape lion 1 | 0.4866698 | 0.5010742 | 0.4722653 | Male | Unknown |
| Cape lion 2 | 0.9152546 | 0.9472044 | 0.8833047 | Female | Unknown |

Layard, E. L. (1861). The catalogue of the south African Museum. *The Cape Argus, Cape Town*.

Methuen, H. H. (1848). *Life in the wilderness: or, wanderings in South Africa*. Richard Bentley.

Pocock, R. I. (1931). The Lions Mane. *The Field, The County Newspaper*, 208.

Sclater, W. L. (1900). *The mammals of South Africa*. RH Porter.

Selous, F. C. (1908). *African Native Notes and Reminiscences*. Macmillan.

Smith, A. (1984). South African Quarterly Journal. 2.

Smith, C. H. (1842). “Black maned lion, Plate X.,” in *The Naturalist’s Library*, ed. W. Jardine (WH Lizars), 177.

Stevenson-Hamilton, J. (1947). Wild life in South Africa. *(No Title)*.

Summers, R. F. H. (1975). *A History of the South African Museum, 1825-1975. Compiled from Museum Records.* , ed. A. A. Balkema Cape Town.

Thom, H. B. (1952). *Journal of Jan van Riebeeck; 1656-1658.* , ed. A. A. Balkema Capetown: For the Van Riebeeck Society.
